# ELViM: Exploring Biomolecular Energy Landscapes through Multidimensional Visualization

**DOI:** 10.1101/2024.01.04.574173

**Authors:** Rafael G. Viegas, Ingrid B. S. Martins, Murilo N. Sanches, Antonio B. Oliveira, Juliana B. de Camargo, Fernando V. Paulovich, Vitor B.P. Leite

## Abstract

Molecular dynamics (MD) simulations provide a powerful means to explore the dynamic behavior of biomolecular systems at the atomic level. However, analyzing the vast datasets generated by MD simulations poses significant challenges. This manuscript discusses the Energy Landscape Visualization Method (ELViM), a multidimensional reduction technique inspired by energy landscape theory. ELViM transcends one-dimensional representations, offering a comprehensive analysis of the effective conformational phase space without the need for predefined reaction coordinates. We apply ELViM to study the folding landscape of the antimicrobial peptide Polybia-MP1, showcasing its versatility in capturing complex biomolecular dynamics. Using dissimilarity matrices and a force-scheme approach, ELViM provides intuitive visualizations, revealing structural correlations, and local conformational signatures. The method is demonstrated to be adaptable, robust, and applicable to various biomolecular systems.

## Introduction

In recent decades, molecular dynamics (MD) simulations have emerged as a potent tool for unraveling molecular systems’ intricate kinetics and dynamics at the atomic level.^1^ These simulations owe their robustness to significant strides in computational capabilities, modeling techniques, and sampling methods.^2–5^ These advancements have extended simulation timescales and enabled the modeling of larger and more complex systems. However, analyzing the extensive datasets generated by MD simulations remains a formidable challenge. Researchers often rely on identifying meaningful reaction coordinates or collective variables to extract biophysically relevant information from molecular trajectories. Moreover, in recent years, machine learning techniques have also emerged as valuable tools for analyzing molecular trajectories.^6–8^

Analyzing molecular trajectories encounters several difficulties due to the large number of degrees of freedom, leading to a high-dimensional phase space. Each configuration in an MD trajectory can be conceptualized as a vector in a high-dimensional space. To illustrate, consider the description of the configurational space using only the alpha carbons of a protein with N residues; each conformation then exists as a vector within a multidimensional space encompassing 3N dimensions. Extracting meaningful features from this multidimensional space becomes arduous due to the so-called “curse of dimensionality”. In such high-dimensional spaces, data points representing protein configurations are notably sparse, making fundamental analyses such as clustering exceptionally challenging.^6,9^

Various strategies have been employed to tackle these challenges, with dimensionality reduction techniques (DR) standing out. These methods aim to reduce the dimensionality of the configurational space, enabling more manageable analysis and visualization. The difficulty relies on finding a good set of reaction coordinates. Another approach involves using DR techniques to map the multidimensional space onto a more manageable, lower-dimensional space. When the lower-dimensional space comprises only two or three dimensions, it facilitates the visualization of an MD trajectory using scatter maps, where each sampled conformation is represented by a data point. The underlying assumption is that the lower-dimensional representation can eliminate noise and redundant information while preserving only the relevant features of the multidimensional space.^6,8^

The first DR technique applied to MD trajectories was the Principal Component Analysis (PCA),^10–12^ a linear technique that works by diagonalizing a covariance matrix and projecting the data along the eigenvectors that retain the largest data variance, and it has been widely applied to several biological and chemical systems. Multidimensional Scaling (MDS) ^13–16^ is a set of methods that aim to preserve pairwise distances or dissimilarities estimated in the high-dimensional space in the low-dimensional space. MDS methods differ in how distances are estimated and how the optimization is carried out to represent these distances on the plane. However, both PCA and MDS are linear methodologies that rely on the hypothesis that most of the variance in the multidimensional data can be captured by a hyperplane.^6^ Addressing limitations of these linear methods, recent advancements have introduced nonlinear methods such as kernel PCA,^17^ Diffusion maps,^18^ Isomap,^19^ t-SNE,^20^ UMAP,^21^ Sketch-map,^22^ and EncoderMap.^23^ While some methods seek a mapping between the high and low-dimensional spaces, others are developed to visualize the data by optimizing the positions of points in the low-dimensional space to satisfy some cost function.

Another approach, inspired by statistical mechanics and spin glass systems principles, is the energy landscape theory (ELT),^24^ which has been primarily used to describe the folding process. This theory depicts a funnel-shaped free-energy landscape biased towards the native state of a protein. In this representation, an ensemble of unfolded structures populates the high-energy portion of the landscape, which funnels toward the native structure. While this approach is not restricted to studying folding, it applies to a wide range of biomolecular systems and functions. Even though these processes are intrinsically multidimensional, it is often feasible to describe the kinetic and thermodynamic properties in terms of a few key quantities. However, one limitation of these techniques is that they require predefined reaction coordinates, potentially masking the richness and details of the problem.

This manuscript elaborates on the Energy Landscape Visualization Method (ELViM), inspired by the ELT. Initially devised for visualizing the protein folding funnel,^25,26^ ELViM represents a significant advancement by transcending the one-dimensional representation. ELViM is a multidimensional reduction method based on internal distances between pairs of structural conformations of the entire analyzed dataset. Moreover, the method does not depend on reference structures or any other reaction coordinate. Through an iterative process, the method seeks to project an ensemble of conformations into two optimal dimensions, facilitating an intuitive visual analysis of the energy landscape. ELViM has been successfully applied in the study of various biomolecular systems, including an RNA tetraloop,^27^ ordered proteins,^26,28–30^ and intrinsically disordered peptides^31,32^ and proteins.^33^ We begin by discussing the general aspects of the method, subsequently delving into its intricacies, such as the dissimilarity metric, code details, iteration steps, and auxiliary tools, which are available at github (https://github.com/VLeiteGroup/ELViM). We illustrate the method using the folding of the MP1 peptide and deliberate on ELViM’s potential and caveats.

## Methods

### ELViM

Given an ensemble of conformations, the analysis using ELViM comprises two fundamental steps for generating lower-dimensional visualizations of the data set:

- Dissimilarity Matrix Calculation: In the first step, ELViM calculates a dissimilarity matrix. This matrix contains estimates of pairwise structural distances between conformations in the data set.
- Multidimensional Projection: In this process, each conformation from the molecular trajectory is represented as a data point in a two-dimensional space. This lower-dimensional representation is referred to as the “effective phase space” or simply the “ELViM projection”.

In the upcoming sections, we provide detailed information regarding these main steps.

### Dissimilarity Matrix

The initial implementation of ELViM was for lattice systems,^25^ and a dissimilarity metric based on the ratio between the Jaccard index and the Jaccard distance was introduced. For out-of-lattice biomolecular systems, Oliveira et al. ^26^ proposed a novel dissimilarity metric based on the order parameter *Q_w_*.^34,35^ Considering two conformations, denoted as k and l, the similarity between them is defined as follows

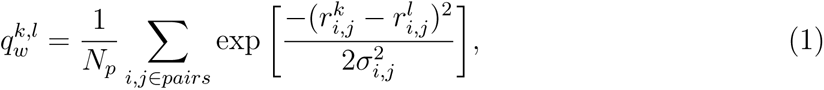

where 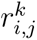 and 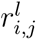 are the distances between the α-carbon of residues i and j from conformations k and l, respectively. *N_p_* is a normalization constant equal to the number of α-carbon pairs, and σ*_i,j_* sets the similarity resolution of the metric and is defined by

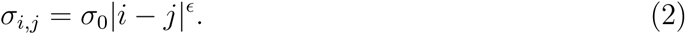

Typically, the values of σ_0_ and ɛ are set to 1 Å and 0.15, respectively. ^35^ However, these parameters can be adjusted to fine-tune the dissimilarity metric for different systems. The dissimilarity between the pair of conformations k and l is defined by

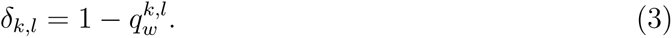

This dissimilarity measure is unitless and falls within the range 0 *≤* δ*_k,l_* < 1. A dissimilarity value equal to zero is only achieved when the two conformations are identical. Notably, this dissimilarity measure only relies on internal distances, not requiring structural alignment. While it is usual to consider only α-carbons for protein systems, this metric can be adapted to incorporate information from all atoms or other coarse-grained representations. After calculating dissimilarity values between every pair of conformations, these computed values are subsequently stored in a dissimilarity matrix used as input by the multidimensional projection technique. When there is a reference conformation (*r*), such as a native state, one can define a reaction coordinate with respect to this structure *Q_w_*, such that 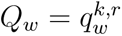 *Q_w_* values can be used to color the data points and provide insights into the overall energy landscape.

### Multidimensional projection

The dissimilarity matrix provides information regarding the structural distances, or dissimilarities, between every pair of conformations. ELViM uses this dissimilarity matrix to perform a multidimensional projection procedure, which aims to project the data from the multidimensional conformational phase space onto a two-dimensional space while preserving, as well as possible, the relevant information. The outcome is a two-dimensional mapping where each data point corresponds to a protein conformation, and the pairwise Euclidean distances in this reduced space are optimized to closely approximate the dissimilarity values calculated within the multidimensional space. This mapping provides accessible and intuitive visualizations of structural correlations among biological macromolecular conformations.

In ELViM, the multidimensional projection optimization is performed using the force scheme technique.^36^ This algorithm, introduced by Tejada et al., is a standard method for multidimensional projection, offering a balance between precision and computational efficiency. It is referred to as a force approach or force-directed method because it treats data points as masses interconnected by springs. This analogy captures how data points are “attracted to” or “repelled by” each other to optimize their pairwise distances to approximate the original dissimilarity. In this algorithm, each conformation k with coordinates 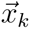 in the multidimensional phase space is projected to a data point 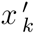, represented in the Cartesian plane by the coordinates 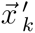. The projection begins with a random arrangement of data points to form the initial projection (X_0_).

In each step, a conformation k acts as a reference, and the projected positions of all other conformations *l* (*l* ≠ *k*) are slightly perturbed to adjust the distance in the plane 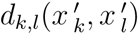 with the dissimilarity estimated in the multidimensional phase space δ*_k,l_*(*x_k_*, *x_l_*). This adjustment is always carried out in the direction of the vector 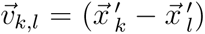 and is proportional to the difference between the dissimilarity and the Euclidean distance (δ*_k,l_−*d*_k,l_*). Using a gradient-descent like method, the cost function to be minimized is given by

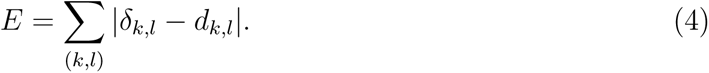

The degree of perturbation is controlled by a learning rate parameter, *L_r_*, which may vary from an initial value to a predetermined minimum value *L_r_min__*, both specified by the user. The learning rate for the i-th iteration is set to be

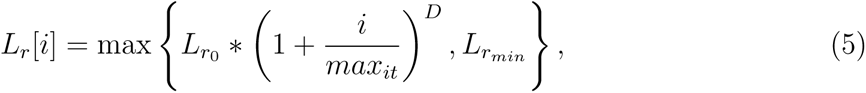

where *L_r_*_0_ is the initial learning rate value, *D* is the decay exponent and *L_r_min__* is the minimum learning rate value. Note that combining the maximum function and the parameter *L_r_min__* allows the combination of an initial annealing of the learning rate followed by constant learning rate steps. Typical value ranges are: 1/8 < *L_r_*_0_ < 0.5, 0.95 < D < 3, and 0 < *L_r_min__* < 1/8.

The algorithm as used in ELViM is detailed as follows:

#### Algorithm 1 ELViM-Force Scheme Algorithm

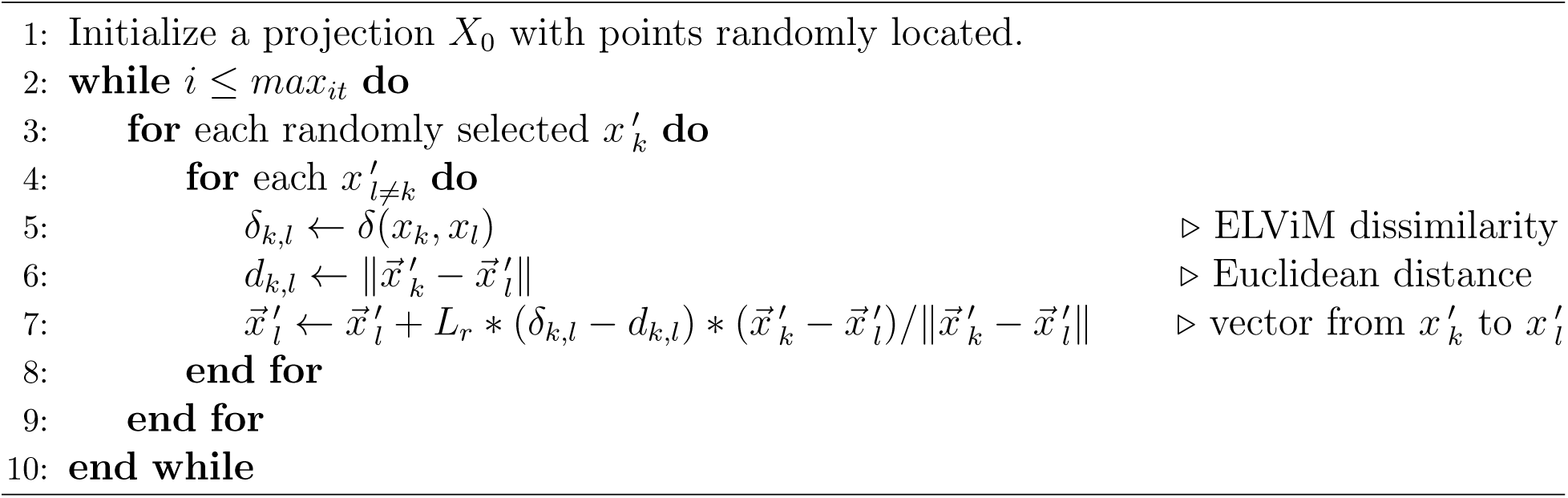

### ELViM code details

The ELViM main program is implemented in Python 3. It is based on the force scheme routine authored by F.V. Paulovich (https://github.com/fpaulovich/dimensionality-reduction). To handle molecular trajectories and extract the *C_α_* coordinates for evaluating the dissimilarity matrix, the program utilizes MDtraj,^37^ a Python library for molecular dynamics analysis. Additionally, the program benefits from parallel processing on CPU cores using Numba,^38^ which optimizes Python code and executes it efficiently on CPU hardware.

ELViM accepts as input molecular trajectory files in various formats recognized by MD-Traj, including single trajectory files (e.g., PDB) or binary trajectories (e.g., XTC, DCD) along with a reference topology file (e.g., PDB). ELViM also offers the option to save the calculated dissimilarity matrix in a Python binary format for future use with different projection parameters. Alternatively, it allows the use of precomputed dissimilarity matrices. The program’s output consists of a text file containing Cartesian coordinates representing the position of the biomolecular conformations within the effective phase space.

### Simulation Details

To illustrate the utility of ELViM in analyzing the structural properties of a biomolecular system, we conducted the analysis with the antimicrobial peptide Polybia-MP1 (or MP1).^39^ The MP1 peptide was constructed on alpha-helix configuration on VMD plugin Molefacture, solvated in a water box measuring 55×55×55 Å, containing NaCl ions in order to neutralize the system and also have 150 mM salt concentration. The system was equilibrated with 10000 steps of conjugate gradient energy minimization and 10 ns of equilibrium MD with backbone restraint. The simulations were performed in the NPT ensemble, at 330 K, with a 2 fs time-step and periodic boundary conditions, using the software NAMD.^40^ Temperature and pressure were modulated by Langevin thermostat^41^ and Langevin piston, ^42^ respectively. The SHAKE algorithm^43^ was used to constrain the lengths of covalent bonds, and the geometry of water molecules was preserved using the SETTLE algorithm. ^44^ The Van der Waals interactions were calculated with a cut-off of 12 Å with a switching distance of 10 Å, and the long-range interaction was treated using the Particle Mesh Ewald (PME) method.^45^

## Results

In this example, we employed the ELViM to project an effective conformational phase space of the MP1 peptide sampled from an all-atoms MD simulation. Along 600 ns of MD trajectory, we observed folding/unfolding events, when the peptide structure fluctuates between α-helix and random coil conformations. Figure 1.a displays the time evolution of the reaction coordinate, *Q_w_* and the RMSD, both calculated having as a reference the conformation that minimizes the energy of the α-helical structure. In this case, *Q_w_* equals 1 for an ideal α-helix and reached 0.2 for unfolded conformations.

**Figure 1:**
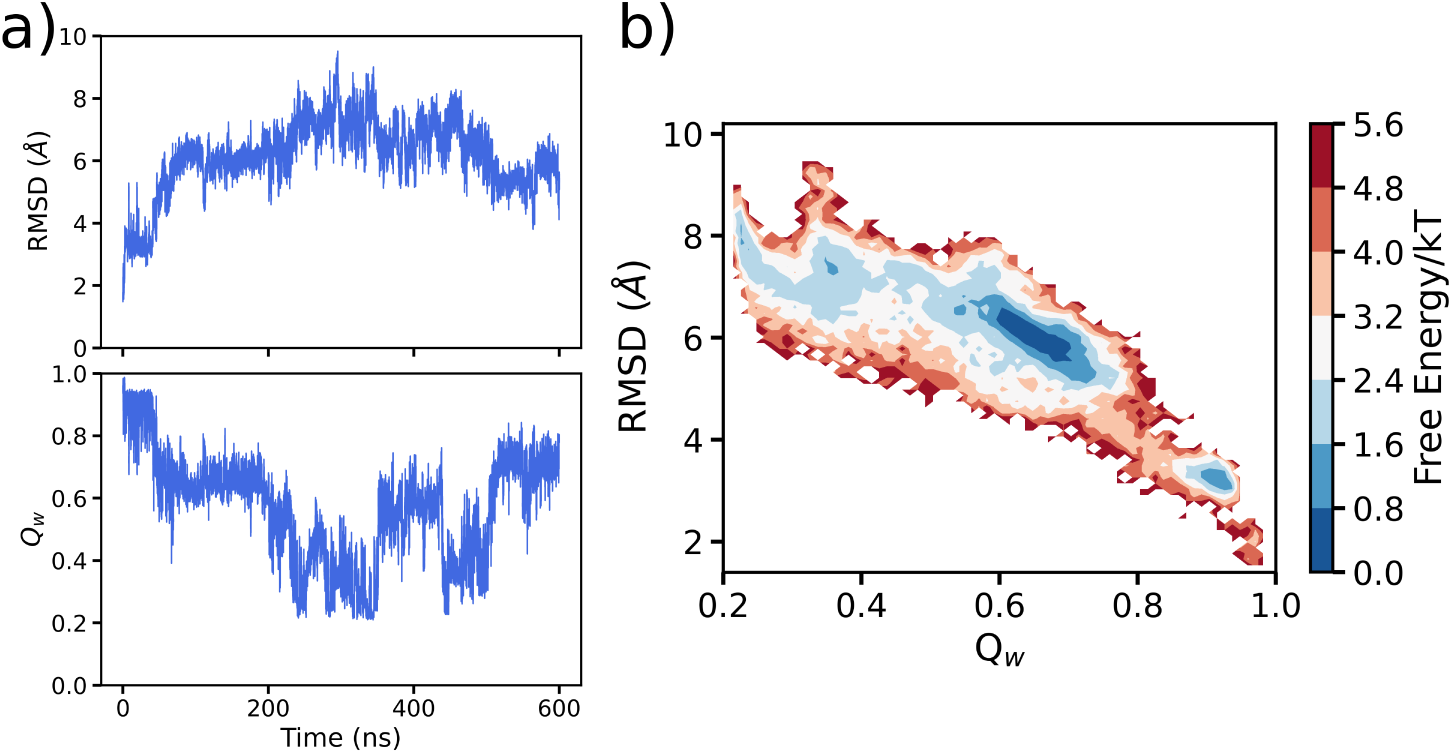
MD Trajectory. (a) Reaction coordinate *Q_w_*, and RMSD were both calculated using *C_α_* coordinates and taking as reference the conformation that minimizes the energy of the α-helical structure. (c) 2D projection of the free energy landscape as a function of the RMSD and *Q_w_*.

A common approach to comprehensively describe the sampled conformational phase space involves projecting the free energy landscape onto a plane defined by order parameters or reaction coordinates of biological significance. In Figure 1.b, a free energy estimation is presented through the projection of RMSD and *Q_w_*. Analysis of this surface reveals a predominant basin of partially-folded conformations (with Q *≈* 0.65), accompanied by two smaller basins: one representing the folded state, characterized by high values of *Q_w_*and low RMSD values, and the other representing the unfolded state, characterized by low *Q_w_* values and high RMSD values. Although this analysis is valuable, particularly in tasks such as estimating free energy barriers and discerning intermediate or metastable states, it does have limitations. Notably, this analysis does not offer a detailed structural mapping, primarily due to the degeneracy associated with these order parameters, which would provide atomistic insights into molecular mechanisms.

The results of applying ELViM to analyze the MP1 MD trajectory are depicted in Figure 2. Dissimilarity was computed using a σ_0_ value of 1 Å. The learning rate varied from 0.3 to 0 with a decay of 0.95. In this representation of the landscape, each dot corresponds to a conformation. The axes have been omitted to enhance clarity, as pairwise distances are the only meaningful data. In Figure 2a, the dots are color-coded based on their *Q_w_* values, and we provide four representative structures. In this depiction, unfolded conformations are colored in dark blue, while the folded conformations are colored in dark red. By coloring the dots according to various biophysically relevant variables, we can obtain different insights into the system. As additional examples, we have also colored the ELViM projection based on the radius of gyration *R_g_* (Figure 2.b) and the content of α-helix (Figure 2.c).

**Figure 2:**
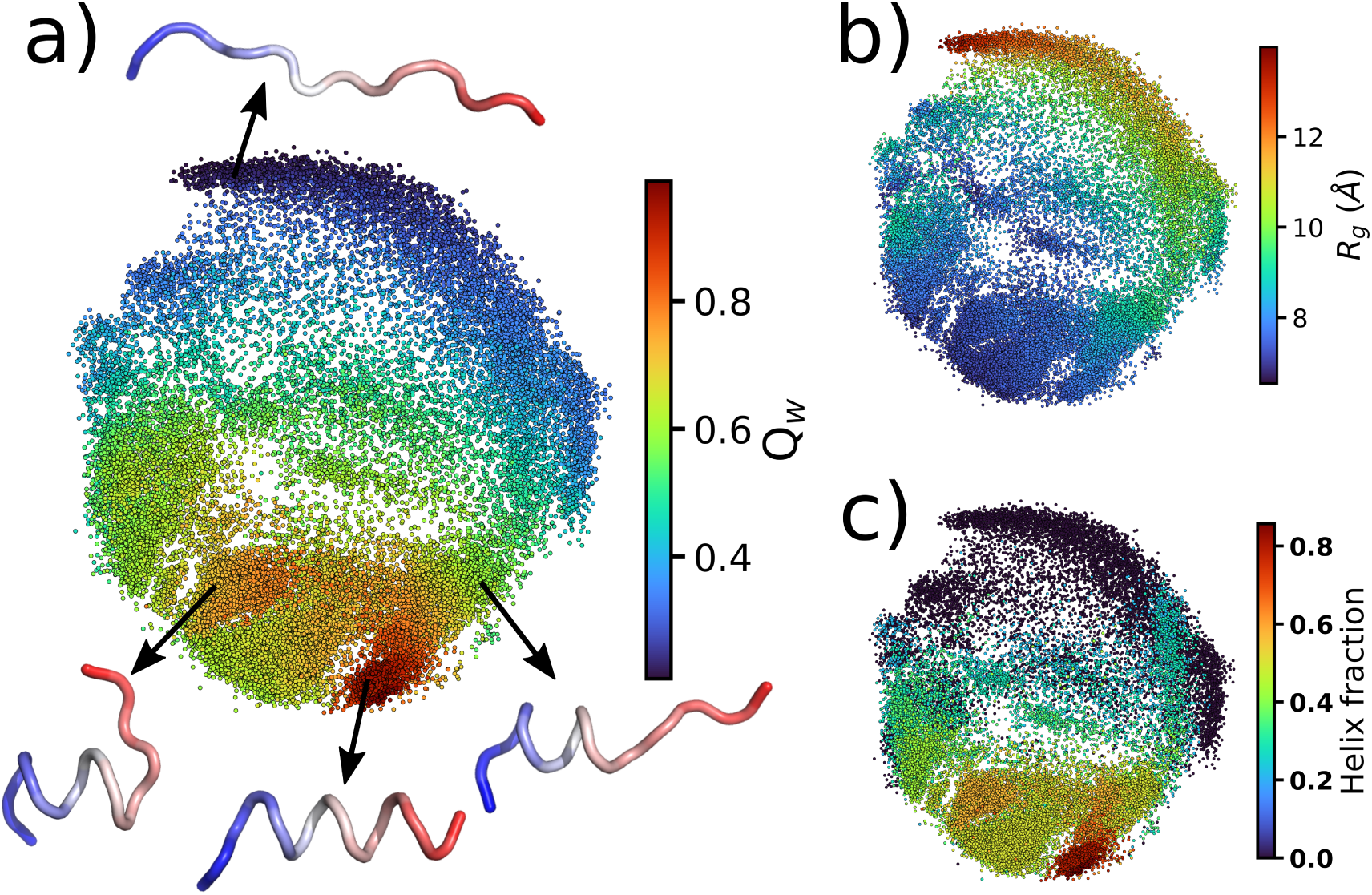
ELViM projection. Each conformation is represented by a dot. The x and y-axis were omitted as they have no meaning. A heatmap is employed to show the values of (a) the reaction coordinate *Q_w_*, (b) the radius of gyration (*R_g_*), and (c) the helix content percentage for each conformation. Some conformations were selected from regions indicated by arrows and shown in carton with N-terminus in blue.

To gain further insights into the folding landscape, it is essential to analyze how the density of data points varies throughout the effective conformational phase space. This measure estimates the density of states in the ELViM projection and allows for identifying basins formed by similar structures frequently visited during the simulation. Here, density is calculated using a Gaussian Kernel Density Estimate (KDE) method, as implemented in scipy.^46^ The results are depicted in Figure 3, where we label some of the higher-density regions from (i) to (viii).

**Figure 3:**
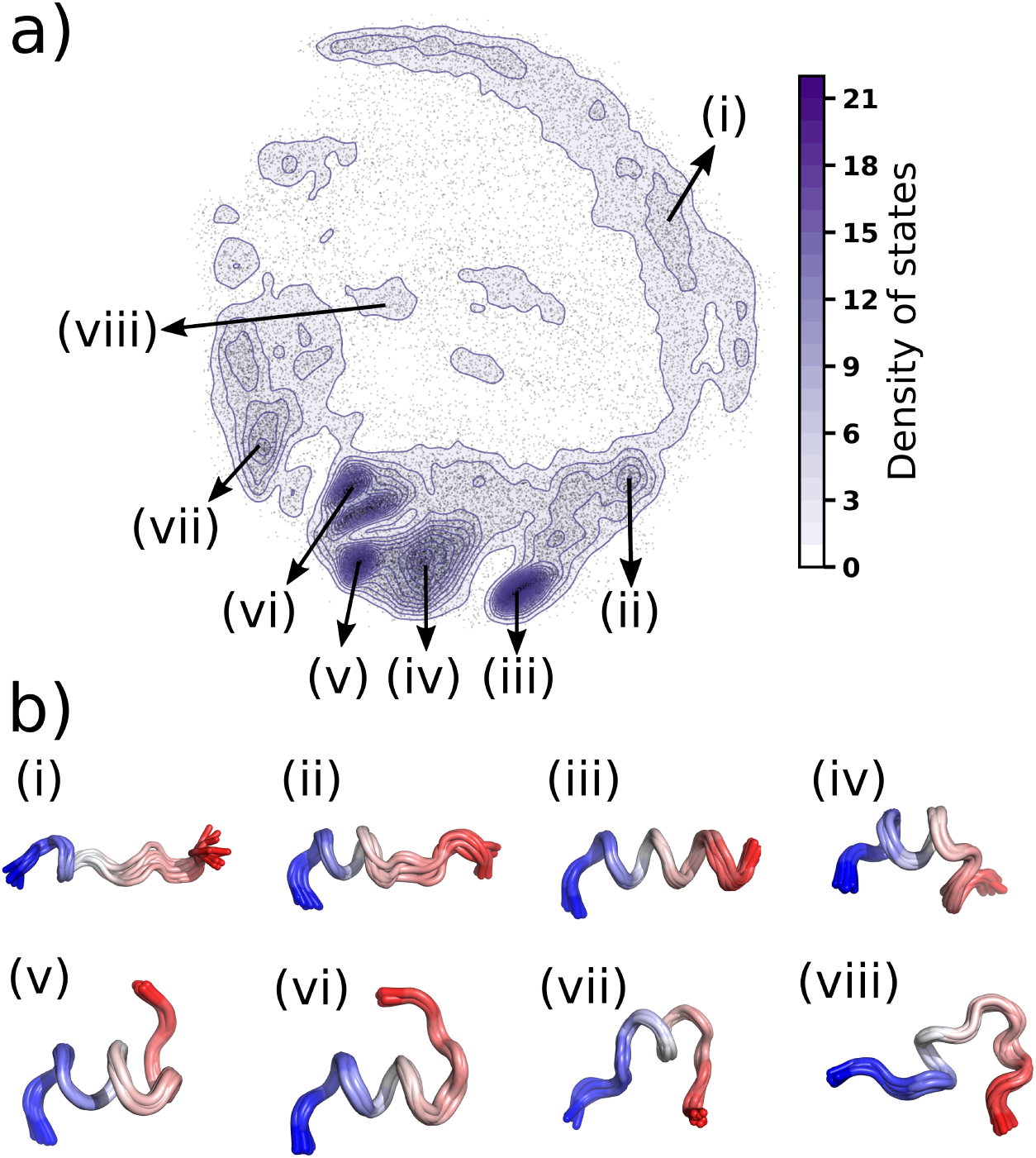
Density of states and Local Conformational Signature (LCS). (a) The density of states is calculated using a Gaussian Kernel (KDE). The density is indicated by the colormap and contour plot superimposed to the ELViM projection dots in gray. (b) Data points from eight regions were manually selected and 10 representative structures (LCS) of each region are depicted superimposed.

Figure 3b presents representative structures from these selected regions, referred to as Local Conformational Signatures (LCS). To identify an LCS, we manually select an arbitrary region and calculate a matrix of distance-RMSD (dRMSD) ^47^ values to find a reference conformation, which is defined as the conformation that minimizes the average dRMSD. The LCS displays this conformation superimposed to its n closest neighbors according to dRMSD values. It is noteworthy that this method is not a clustering analysis; rather, it is a visualization tool designed to provide a structural signature of conformations from arbitrarily selected regions. The representative structures from all high-density regions grant a broader perspective of the effective phase space. Furthermore, this approach allows us to compute contact maps and other biophysically relevant variables for conformations in an LCS. In cases where the data originates from unbiased simulations, high-density regions likely indicate local free energy minima within the overall free energy landscape. Consequently, this analysis can provide essential insights, revealing atomistic details regarding these basins.

## Discussion

The ELViM algorithm starts with a random initialization of the projection and then optimizes distances to minimize the cost function (Eq. 4). To prevent dependencies on iteration order, points are randomly selected during optimization. Consequently, different runs produce different projections for the same set of parameters. If the complexity of the system is not too high, the projection stabilizes, and the different run outcomes are slightly different from each other, but the global structure of the effective phase space is maintained. As an example, we have generated three independent projections for the effective space of MP1 and compared them in Figure S1 of supporting information. As previously discussed, only pairwise distances are meaningful in ELViM projection, and rotations and reflections about arbitrary axes can be performed without changing the global structure of effective phase space. In Figure S1, we also arbitrarily selected seven local groups and showed how their position and composition are maintained throughout different replicas of the projection.

Based on our experience, a number of iterations equal to the square root of the number of conformations used in the projection typically suffices to achieve convergence. However, complex systems may require a larger number of iterations. To ensure the convergence of the projection with consistent global features, it is advisable to run and compare multiple projections. Consistency in the relative positioning of main basins is also a good indicative of convergence.

In the ELViM implementation, certain parameters can be adjusted to fine-tune the projection results. For instance, users can specify the parameters in the σ*_i,j_*(Eq. 2), which determines the dissimilarity resolution. While ELViM projections are usually robust when using the typical σ*_i,j_* parameter values indicated in the methodology section, extreme values may distort the projection. A small value tends to excessively increase the dissimilarity between data pairs, making it challenging to represent all the data points onto the plane. Conversely, a value that is too large may result in lower dissimilarity for very different structures, causing them to be placed within neighboring regions of the projection. To illustrate this dependency, we provide in supporting information two ELViM projections for the MP1 system using different σ_0_ parameters (Fig. S2).

Initially, our implementation considered only *C_α_* carbons for dissimilarity metrics. However, the code can be easily adapted to include other representations, such as using distances between all heavy atoms, as demonstrated in our RNA tetraloop study.^27^ When atom indices differ from residue indices, adjustments to the σ*_i,j_* definition may be necessary. Modifications to the cost function (Eq. 4) can also be explored, such as squaring the residuals’ differences between dissimilarities and projected distances in Eq.4. However, these modifications should be carefully evaluated for each specific system studied.

The learning rate parameter (Eq. 5) comprises three adjustable constants: the initial value *L_r_*_0_, final value *L_r_min__*, and an exponent controlling the decay rate D, as discussed in the methods section. This parameter acts as a scaling factor, which multiplies the residual difference between dissimilarities and distances, setting the size of the perturbation applied to every point in each iteration step. In this way, these parameters influence both convergence and the final projection outcome. Using excessively small learning rate values may confine data points within incorrect neighborhoods and significantly increase the required number of iterations to achieve convergence. Conversely, an excessively high learning rate may lead to an unstable projection. Large values of *L_r_min__* may result in isolated structures resembling islands if they have considerable dissimilarity to their nearest neighbors. An example of such behavior is provided in supporting information Fig S3.

For projections with sufficient statistics, the probability density of each region in the projection phase space is associated with its free energy. Therefore, ELViM can offer an intuitive representation of the free energy landscape as sampled during a simulation. The density of data points in the projection can be estimated using a two-dimensional histogram or kernel density estimation (KDE), which can then be used to calculate free energy differences. In cases where sampled conformations originate from enhanced sampling simulations or different conditions, data points may no longer carry the same weight. In such instances, ELViM can still provide a useful representation of the configurational space for visualization, but calculating free energies may necessitate a reweighting procedure. For detailed examples, refer to our previous works.^32,48^

Finally, it should be acknowledge the occurrence of projection errors or distortions in dimensionality reduction and multidimensional visualization techniques. These distortions are caused by data points misplaced in the projection.^9,49^ These errors may be due to inherent mathematical limitations of projecting high-dimentional data onto a two-dimensional space.^9^ In this context, some quality metrics have been proposed to quantify the extent to which distances or neighborhood relationships are preserved in the projection.^50,51^ However, it is important to note that these metrics primarily assess the effectiveness of the optimization procedure but do not directly measure the method’s capability to represent the unknown topology of the multidimensional phase space. ^8^

When analyzing the ELViM projection, misplaced points would behave as structural noise in a narrow neighborhood. In this sense, we suggest that the projection may be interpreted based on the prevalent structure signatures within a local neighborhood. We provide the LCSs tool to address this purpose, which identifies prevalent conformations within an arbitrarily selected region. By analyzing various LCS comprising all the high-density regions of the projection, it is possible to picture how representative conformational states are distributed throughout the projection. As previously stated, this tool provides a qualitative analysis aiming at an intuitive interpretation of the projection. However, maintaining awareness of misplaced data points remains important to ensure the quality of the projection.

One intrinsic limitation of ELViM is associated with the size and complexity of the investigated system. From a mathematical standpoint, multidimensional projection can be considered as an ill-posed problem. The projection iteration procedure may converge to qualitatively different degenerate solutions, which may be viewed as alternate descriptions of the landscape. The limits of applicability of ELViM in terms of the size and complexity of a studied system have yet to be tackled and will be addressed in future studies.

### Conclusions

In conclusion, ELViM emerges as a valuable tool for unraveling the intricacies of biomolecular energy landscapes. Its ability to navigate high-dimensional configurational spaces and provide intuitive visualizations makes it a versatile choice for analyzing MD trajectories and structural ensembles. By eliminating the reliance on predefined reaction coordinates, ELViM allows for a more comprehensive exploration of the conformational phase space. Applying ELViM to the folding landscape of Polybia-MP1 demonstrates its efficacy in capturing dynamic transitions and revealing structural nuances. With its adaptable parameters and robust optimization scheme, ELViM stands as a promising method for researchers seeking a deeper understanding of biomolecular dynamics and functional mechanisms.

## Acknowledgements

The authors acknowledge financial support from Prope/Unesp and the Brazilian agencies FAPESP (Grants 2023/02219-1, 2022/08738-8, 2023/08101-2, and 2021/15028-4) and National Council for Scientific and Technological Development – CNPq (Grant 310017/2020-3). ABOJ acknowledges the Robert A. Welch Postdoctoral Fellow program (grant C-1792). This research was supported by resources supplied by the National Laboratory for Scientific Computing (LNCC/MCTI, Brazil) the SDumont supercomputer, URL: http://sdumont.lncc.br, the Center for Scientific Computing (NCC/GridUNESP) of the São Paulo State University (UNESP) and the “Centro Nacional de Processamento de Alto Desempenho em São Paulo (CENAPAD-SP).

## Notes

### Competing Interest Statement

The authors have declared no competing interest.

