## Supporting Information for "ELViM: Exploring Biomolecular Energy Landscapes through Multidimensional Visualization"

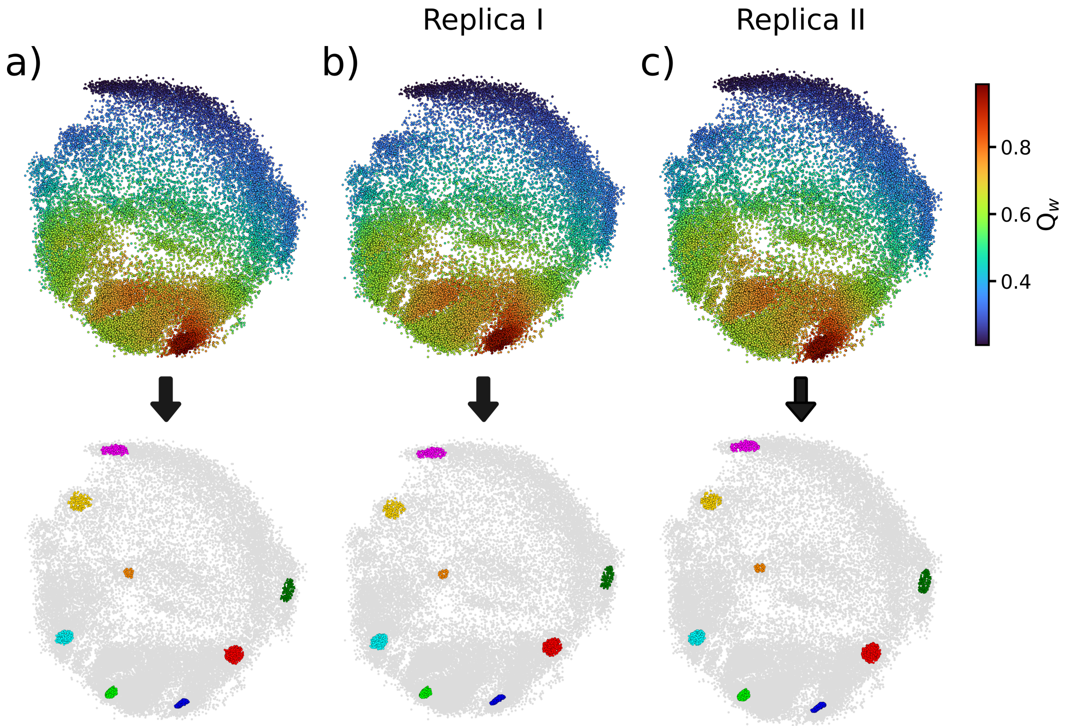


Figure S1: ELViM Reproducibility. The ELViM optimization procedure is inherently stochastic, resulting in slight variations in outcomes across different runs. These observed discrepancies might depend on the complexity of the system. To investigate reproducibility, we replicated the ELViM projection from the main text (a) and generated two independent replicas (b) and (c), all using identical parameter sets. Since the x- and y-axis have no particular meaning, we aligned the ELViM projections for comparison through rotations and reflections about arbitrary axes. Additionally, to illustrate how local and global neighborhoods are preserved across different runs, we selected eight groups of conformations arbitrarily. These groups were consistently represented with the same colors across all replicas (lower panels). Notably, data points within these selected groups consistently maintained cohesion and retained the same relative positions within the projection. This observation suggests that the ELViM projection has converged, emphasizing the robustness of the methodology.

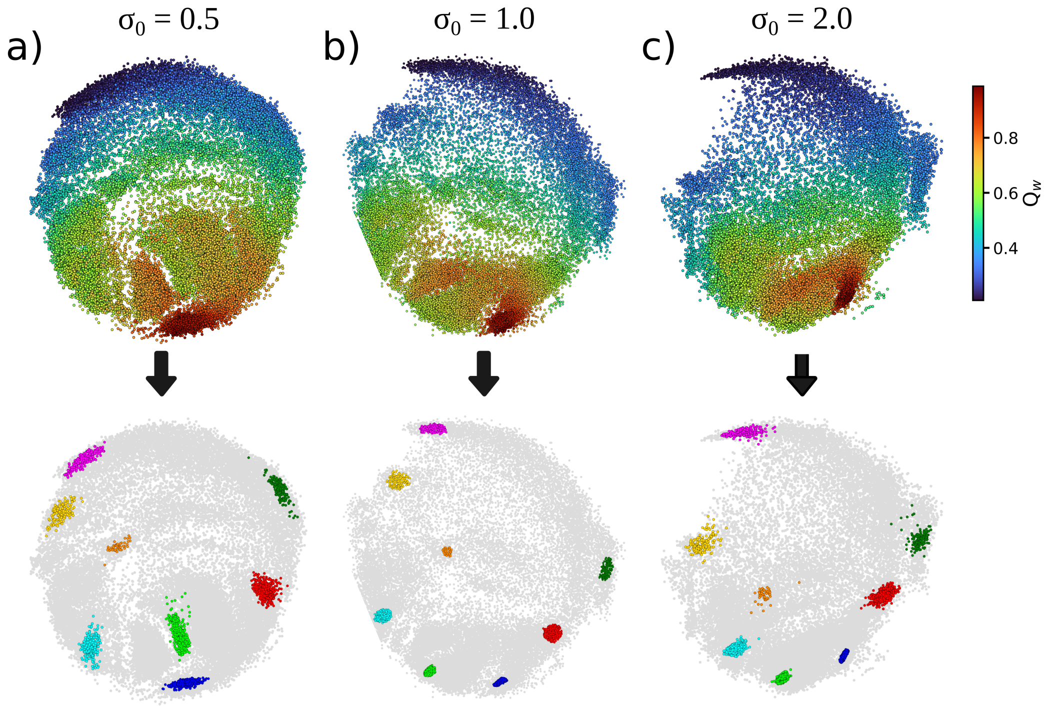


Figure S2: Effect of the parameter $\sigma_0$ on the dissimilarity matrix and ELViM projection. The $\sigma_0$ parameter sets the dissimilarity resolution, wherein larger $\sigma_0$ values lead to smaller dissimilarities between conformation pairs. To illustrate this, we compare three ELViM projections using (a) $\sigma_0 = 0.5$, (b) $\sigma_0 = 1$, and (c) $\sigma_0 = 2$. All projections utilized the same learning rate parameter ($L_{r_0} = 0.3$, $D = 0.95$, $L_{r_{min}} = 0$). Lower values of $\sigma_0$, as seen in projection (a), increase pairwise dissimilarities, potentially revealing more details within folded regions, where dissimilarities are typically small. However, this may lead to excessively large dissimilarities within the unfolded basin, causing projection instabilities, especially for more complex systems. On the other hand, higher values of $\sigma_0$, as illustrated in projection (c), decrease pairwise dissimilarities, potentially collapsing some projection regions. Nevertheless, both local and global information are qualitatively preserved across the three projections, as evident when analyzing the locations of arbitrarily selected groups (lower panels, same groups as in Fig. .

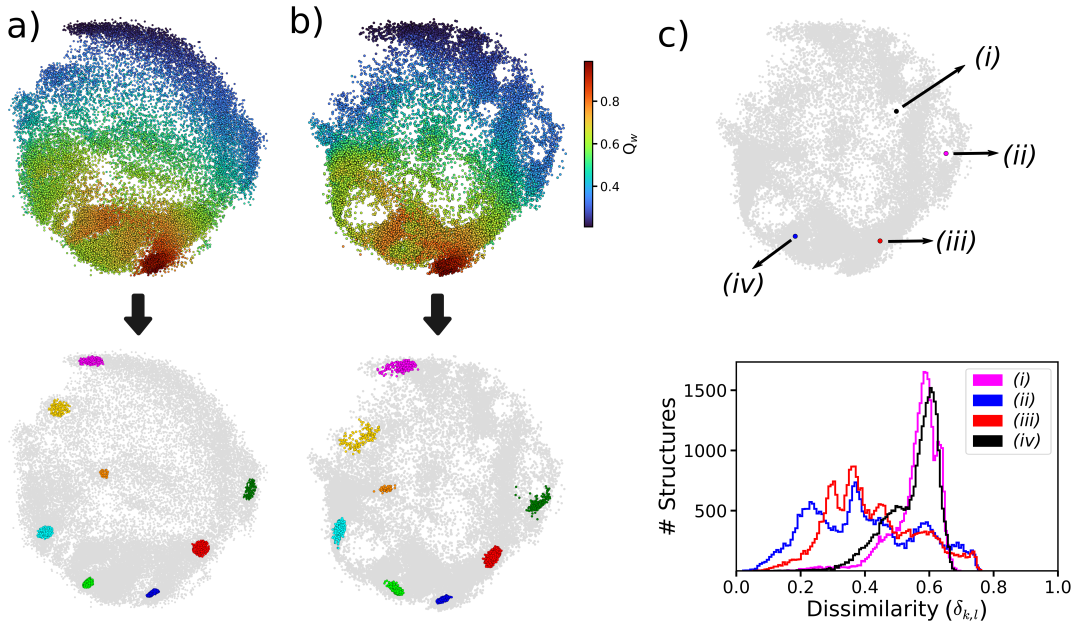


Figure S3: Effect of the learning rate parameter $L_r$ on the ELViM projection. The total $L_r$ represents the fraction of the residual error ($|\delta_{k,l} - d_{k,l}|$) by which a data point is moved during each optimization step. The learning rate is comprised of three free parameters: the initial learning rate ($L_{r_0}$), decay ($D$), and minimum learning rate ($L_{r_{min}}$). To illustrate, we compare two cases: (a) an annealing regime with ($L_{r_0} = 0.3$, $D = 0.95$, $L_{r_{min}} = 0$), as used in the main text, and (b) a constant learning rate with ($L_{r_0} = 1/8$, $D = 0$, $L_{r_{min}} = 1/8$). For both cases, $\sigma_0 = 1$. Larger values of $L_{r_{min}}$ lead to isolated structures resembling islands, as observed in the projection in (b), causing slight distortions in some regions. Excessive or overly large island-like structures may result in unstable and less reproducible projections. In both cases presented here, local and global information are qualitatively preserved across the projections, as observed by analyzing the locations of arbitrarily selected groups (lower panels, same groups as in Fig. S1). Panel (c) shows that conformations prone to form islands exhibit a distinct neighborhood profile. For this analysis, we selected, as shown in the upper panel, two island-like conformations (i) and (ii), and two conformations from high-density regions (iii) and (iv) as reference. The lower panel shows the histogram of the dissimilarity values to all other conformations. Notably, conformations prone to forming islands have fewer close neighbors (dissimilarity $< 0.2$) than conformations from high-density regions.
